# Ultrack: pushing the limits of cell tracking across biological scales

**DOI:** 10.1101/2024.09.02.610652

**Authors:** Jordão Bragantini, Ilan Theodoro, Xiang Zhao, Teun A.P.M. Huijben, Eduardo Hirata-Miyasaki, Shruthi VijayKumar, Akilandeswari Balasubramanian, Tiger Lao, Richa Agrawal, Sheng Xiao, Jan Lammerding, Shalin Mehta, Alexandre X. Falcão, Adrian Jacobo, Merlin Lange, Loïc A. Royer

## Abstract

Tracking live cells across 2D, 3D, and multi-channel time-lapse recordings is crucial for understanding tissue-scale biological processes. Despite advancements in imaging technology, achieving accurate cell tracking remains challenging, particularly in complex and crowded tissues where cell segmentation is often ambiguous. We present Ultrack, a versatile and scalable cell-tracking method that tackles this challenge by considering candidate segmentations derived from multiple algorithms and parameter sets. Ultrack employs temporal consistency to select optimal segments, ensuring robust performance even under segmentation uncertainty. We validate our method on diverse datasets, including terabyte-scale developmental time-lapses of zebrafish, fruit fly, and nematode embryos, as well as multi-color and label-free cellular imaging. We show that Ultrack achieves state-of-the-art performance on the Cell Tracking Challenge and demonstrates superior accuracy in tracking densely packed embryonic cells over extended periods. Moreover, we propose an approach to tracking validation via dual-channel sparse labeling that enables high-fidelity ground truth generation, pushing the boundaries of long-term cell tracking assessment. Our method is freely available as a Python package with Fiji and napari plugins and can be deployed in a high-performance computing environment, facilitating widespread adoption by the research community.

---

Advancements in live imaging technologies^1–7^ have enabled the visualization of cellular dynamics with unprecedented spatiotemporal resolution^8^ over large fields of view^6, 7^ encompassing thousands of cells,^5, 9^ producing vast amounts of multi-dimensional data.^10^ However, one key challenge remains: the accurate reconstruction of cell trajectories and lineages in complex biological systems.^11^ This ability unlocks deeper insights into cellular state and behavior,^9, 12, 13^ tissue mechanics,^14^ morphogenesis,^2, 15, 16^ and regeneration.^17^

Cell segmentation and tracking have been persistent challenges in bioimage analysis.^11^ While cell segmentation has advanced rapidly with the advent of deep-learningbased methods,^18–20^ cell tracking continues to be an open problem.^11^ The primary challenge stems from the accumulation of segmentation errors over time, hindering long-term tracking, especially in dense and dynamic cellular environments. Most automatic tracking methods employ a two-step approach: first segmenting cells, then linking them across time frames.^21–24^ While computationally efficient, this approach struggles with the compounding of errors over time, particularly in dense tissues^25^ or when cells divide rapidly.

Simultaneous segmentation and tracking offers a promising alternative,^26–30^ wherein multiple candidate segments and/or inter-segment links are generated, and then optimal candidates are selected by satisfying biological constraints (*i.e*. cells divide but do not merge) and optimizing specific criteria (*i.e*. only keep links best supported by the data). However, existing methods in this category are constrained by their inability to utilize arbitrary segmentation inputs,^28, 31^ as well as limitations in data scale or dimensionality due to high computational costs.^29, 30^ These factors have limited their applicability to tissue-scale problems.

To address these challenges, we developed Ultrack, a robust and scalable method for large-scale cell tracking that excels even under segmentation uncertainty. Ultrack can track cells (or nuclei) in 2D, 3D, and multichannel datasets, accommodating a wide range of biological contexts. Ultrack seamlessly integrates with various segmentation algorithms, including state-of-the-art deep learning-based cell segmentation tools^18–20, 32–34^ (Fig. 1a). By jointly evaluating candidate segmentations and tracks, Ultrack employs temporal consistency to select the most accurate segments (Fig. 1b). This approach leverages information from adjacent time points to resolve cell segmentation and tracking ambiguities, enhancing performance, particularly in complex, densely packed tissues where cell boundaries are often uncertain (Supp. Video 1). We demonstrate the utility and robustness of Ultrack through a comprehensive set of benchmarks and realworld applications, including top-scoring performance on the Cell Tracking Challenge.^11^ Our method shows improved tracking performance across various use cases and scales, from multi-color and label-free 2D datasets to large-scale 3D time-lapse sequences of developing embryos.

**Fig. 1.**
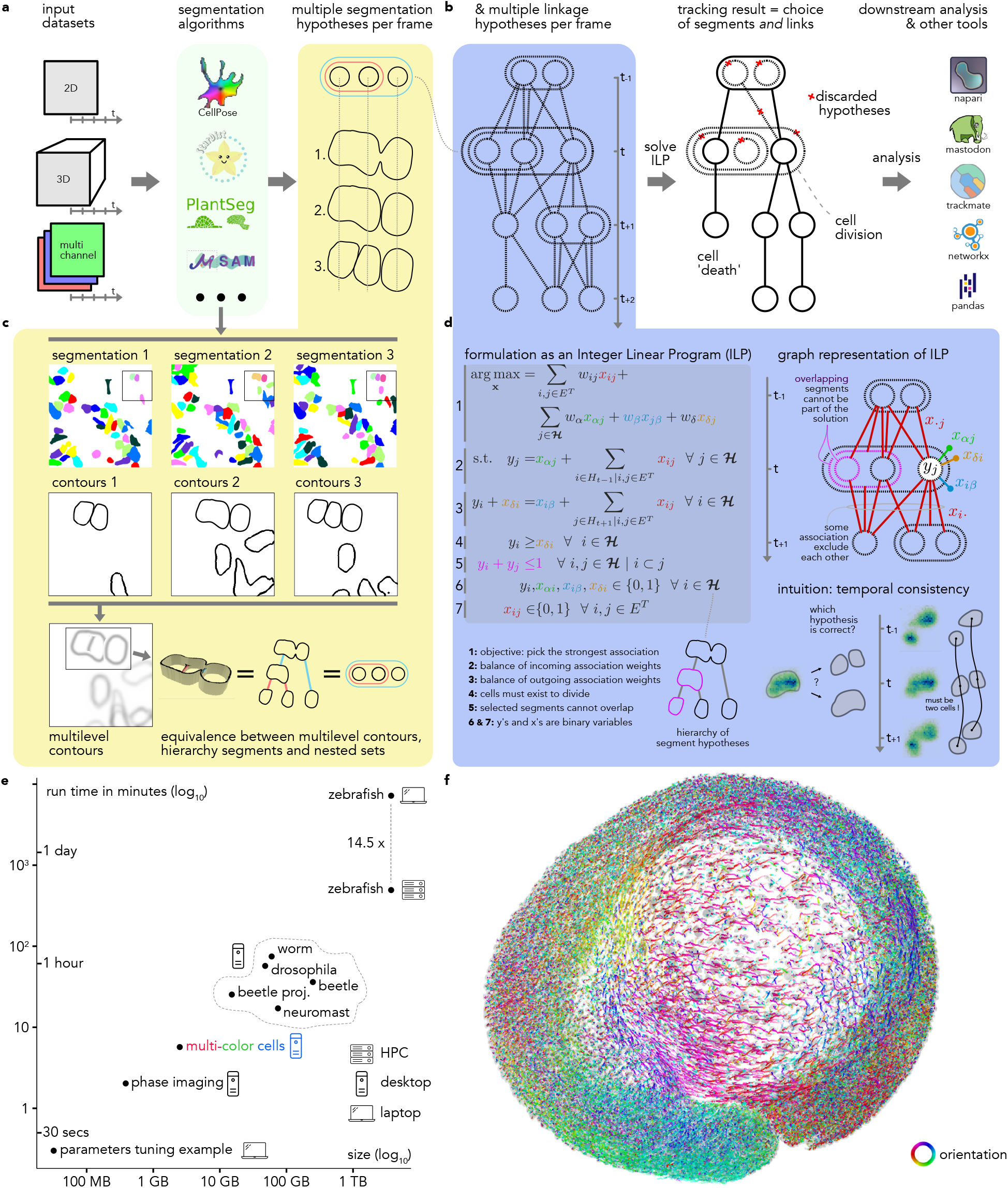
Ultrack overview. **a**, The Ultrack pipeline can process a variety of input images, including 2D, 3D, and multi-color datasets. These images are then processed by any segmentation algorithm or a combination of them. **b** Ultrack builds segmentation hypotheses between frames for tracking and solves an Integer Linear Programming (ILP) problem to identify cell segments and their trajectories. The resulting tracks can be exported in various formats compatible with multiple platforms. **c**, Illustration of how segmentation hypotheses are built using ultrametric contours: multiple segmentations are provided (first row), their binary contours are extracted (second row), and these are combined into a multilevel contour, equivalent to a hierarchy (third row). **d**, Joint segmentation and tracking is performed using an Integer Linear Programming (ILP) formulation, modeling cell behaviors (*e.g*. cell division, cell death, or cells leaving the field of view) while finding non-overlapping cells with maximal association over time. **e**, Ultrack can process arbitrarily sized datasets and scales from a laptop to a High-Performance Computing (HPC) cluster. **f**, Projection of a 3D image of a developing zebrafish embryo, with cell tracks overlaid and colored to indicate track orientation in the xy-plane.

Additionally, we introduce a novel dual-channel sparse labeling approach that enables the generation of highfidelity ground truth for tracking validation. This method combines ubiquitous fluorescence labeling with sparse, random labeling at a distinct wavelength, allowing for the creation of annotations that would be impossible to produce manually. This innovative validation strategy pushes the limits of cell tracking assessment, enabling the evaluation of tracking performance over extended time periods and in increasingly complex tissue environments.

To ensure broad accessibility and ease of use, we provide intuitive interfaces in Fiji,^35^ napari,^36^ Python, and command line, catering to researchers with diverse computational backgrounds. We also made efforts to ensure interoperability with existing popular cell tracking frameworks such as TrackMate^37^ and Mastodon^38^ (see Fig. 1b). Thus, by enabling more accurate, scalable, and accessible cell tracking, Ultrack can accelerate research in developmental biology, cancer research, and regenerative medicine, offering new insights into cellular behavior and tissue dynamics.

## RESULTS

### Avoiding premature mistakes with multiple segmentation hypotheses

Cell tracking accuracy is compromised when cell segmentation (or detection) fails. To address this challenge, Ultrack considers multiple segment hypotheses instead of prematurely committing to a particular choice of segments (see Fig. 1a). These hypotheses can be obtained from the original imaging data pixel intensities, multiple algorithms (*e.g*. watershed plus Cellpose), or from one or several algorithms configured with different parameters. This approach allows the integration of various segmentation methods, maintaining compatibility with the growing array of novel off-the-shelf segmentation models.^39, 40^ However, the main challenge of this approach and previous joint segmentation and tracking methods, as well as the main obstacle for their practical application, is the potentially large number of possible segmentation hypotheses, especially for large multi-dimensional datasets.

### Efficient representation of segmentation hypotheses

To address the challenge of managing multiple segmentation hypotheses, we use ultrametric contour maps (UCMs),^41–43^ which provide a compact representation of these hypotheses. Intuitively, UCMs are *multi-level contour maps* that represent a hierarchy of possible cell boundaries, where the strength of each boundary reflects its likelihood of being a true cell edge (Fig. 1c). This hierarchical structure allows for the efficient encoding of multi-ple possible segmentations within a single map. More formally, UCMs partition space into nested segments, where valid segments do not overlap, or are contained within one another. This property enables Ultrack to efficiently consider multiple segmentation possibilities while tracking cells, even in terabyte-scale datasets (see Fig. 1e), that are challenging for existing joint segmentation and tracking solutions.^26, 28–30^

Hence, the canonical Ultrack input consists of two maps for each time frame: (i) a foreground map distinguishing potential cells from the background and (ii) a grayscale image representing the multilevel contour map (*i.e*., UCM), Fig. 1c. These two input maps can be generated from any method, making Ultrack compatible with a wide range of existing workflows (see Methods for technical details). Therefore, the same algorithm is always applied in every application presented here, and the varying factor is the process of obtaining the multilevel contour map.

### Tracking by maximizing temporal consistency

Once all candidate segments and links between segments are determined, tracking involves solving a complex combinatorial problem of selecting and associating the correct cell segments across time from the set of multiple segmentation and linkage hypotheses (see Fig. 1b and Supp. Video 1). As shown in Fig. 1d, we formulate this combinatorial problem as an Integer Linear Program (ILP)^28, 44^ that simultaneously solves for the optimal selection of temporally consistent segments while adhering to biological constraints. These constraints include cell division, cells entering (*e.g*. moving into the frame), or exiting (*e.g*. cell death or moving out of the field of view), and ensuring that a pixel is not assigned to more than one cell.^45^ This efficient and versatile mathematical formulation leverages highly optimized solvers (Gurobi,^46^ Coin^47^) and thus can handle tens of millions of segments from terabyte-scale datasets (see Fig. 1e). Once solved, the ILP formulation not only selects the most appropriate segments but also encodes the linkage data necessary to reconstruct the cell lineages from the optimal association between the selected segments (see Methods for associations score definition and ILP formulation details). Recapitulating cell lineages and migration patterns, as shown in the developing zebrafish embryo in Fig 1f and Supp. Video Supp. Video 1

### Optimal parameter selection by temporal consistency

Image segmentation techniques, from classical methods like watershed to advanced deep-learning models, face significant challenges in parameter tuning due to several factors: (i) computational: extensive resources are required to explore a large parameter space; (ii) practical: users often lack the expertise to tune parameters effectively; and (iii) technical: a single optimal parameter set may not fit all aspects of a diverse dataset, such as when segmenting cells of varying sizes.

Ultrack addresses these challenges by integrating multiple segmentation labels derived from different parameter settings into a single multilevel contour map. This approach leverages the temporal consistency of the tracking process to select the most appropriate segments for each cell (see Fig. 1d), resulting in more effective segmentation than what could be achieved with any single parameter setting — even the optimal one.

To demonstrate this key feature of Ultrack, we tracked human hepatocarcinoma-derived cells expressing YFPTIA-1^48^ in their nucleus and cytoplasm from the Cell Tracking Challenge (CTC)^11^ using the Cellpose “cyto2” weights^32^ without any fine-tuning. In this dataset, cells have significantly varying intensities, impairing the model’s ability to consistently segment every cell correctly. For example, using original image intensities (*γ* = 1.0), the segmenta-tion model fails to detect dimmer cells (see Fig. 2c – second column from right). Conversely, altering the dynamic range by taking the power of the image intensities with values closer to zero (*γ* = 0.1, 0.25, 0.5) improved the segmentation of dimmer cells but led to incorrect segmentation or non-detection of now-saturated bright cells (Fig. 2c– right most column).

**Fig. 2.**
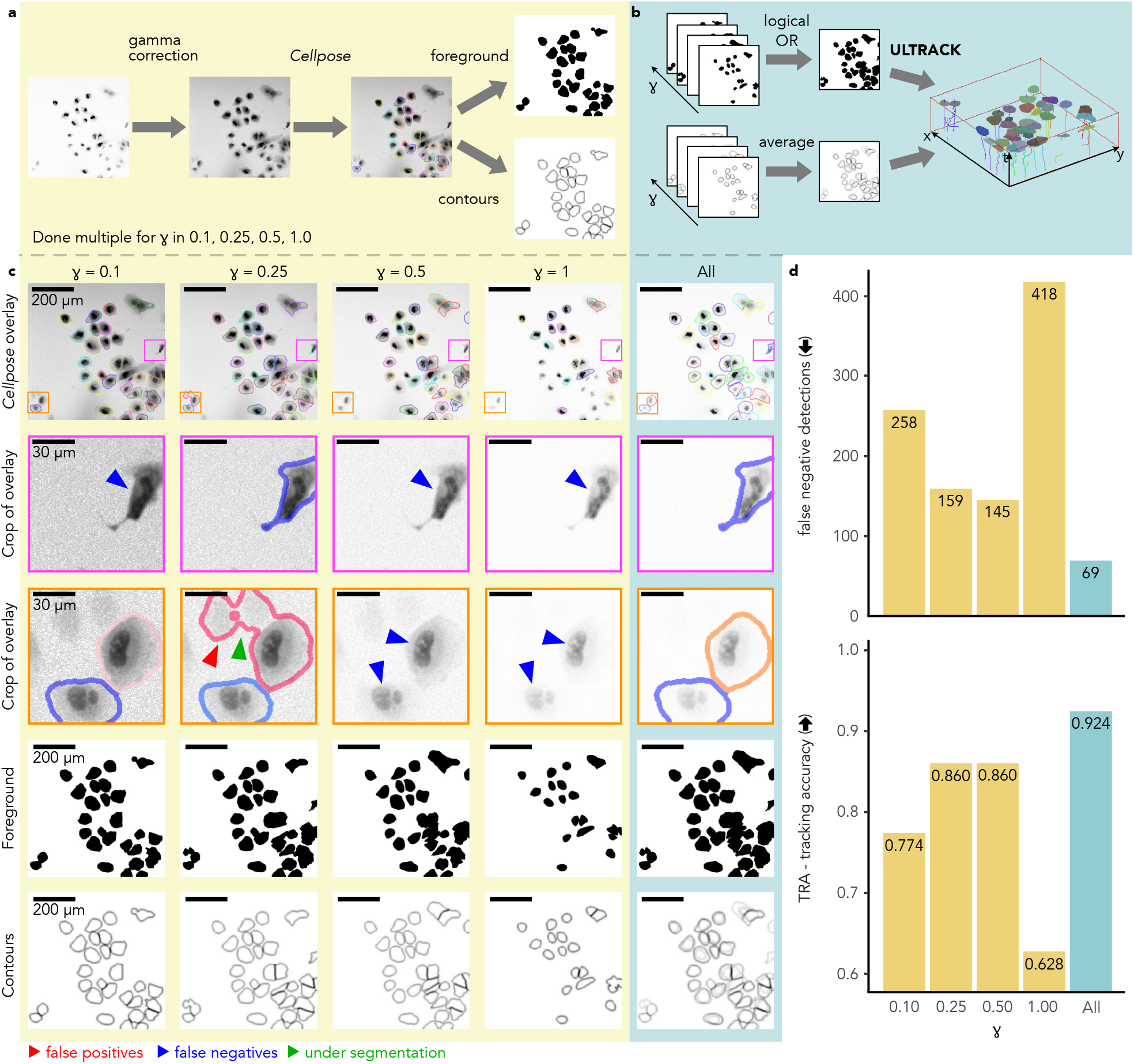
Multiple segmentation hypotheses alleviate the curse of parameter tuning. **a**, Segmentation pipeline using gamma correction with varying settings (*γ* = 0.1, 0.25, 0.5, and 1.0) to generate diverse segmentation hypotheses. **b**, Ultrack’s approach: integrating multiple foreground and contour maps for joint segmentation and tracking. **c**, Impact of gamma correction on segmentation quality: original images (top row), segmentation results (second and third rows) with errors highlighted by colored arrows, foreground masks (fourth row), and contour maps (fifth row). **d**, Quantitative comparison of false negative cell detections and tracking accuracy (TRA) for individual gamma settings and Ultrack’s combined approach.

To segment and track the combination of all parameterizations of gamma, we extracted the binary foreground and contour maps from the Cellpose labels, Fig. 2a. Next, we combined them into a single binary foreground and a multilevel contour map for tracking, Fig. 2b. Further displayed in Supp. Video 2.

We evaluated the tracking and segmentation performance for each individual gamma setting, as well as for the combined contour map approach. To assess performance, we compared the results against the manually curated ground truth provided by the Cell Tracking Challenge (CTC). Figure 2d summarizes the average results obtained from the two publicly available datasets. We used two key metrics for evaluation: the false negative detection rate and the tracking accuracy (TRA).^49^ For false negative detection, lower scores indicate better performance, while for TRA, higher scores are better (see Methods for details).

These results demonstrate that Ultrack achieves more accurate tracking by leveraging multiple segmentation hypotheses, even outperforming the optimal single parameter setting (*γ* = 0.5). This indicates that combining diverse segmentation inputs is often more effective in practice, thus eliminating the need for precise parameter tuning and offering a more robust approach to cell tracking.

### Enhancing cell tracking accuracy through multi-channel integration and color-aware associations

Multi-color labeling has been shown to enhance cell segmentation and tracking capabilities, particularly in complex cellular environments.^50, 51^ However, applying stateof-the-art deep-learning models to such data often requires training on dataset-specific annotations, as pretrained weights are typically optimized for single-channel data. Ultrack addresses this challenge by combining multiple segmentation inputs in two ways: by varying parameters, as demonstrated earlier, and by integrating outputs from different segmentation algorithms applied to separate color channels.

We demonstrate this key feature of Ultrack on a three-channel “multi-color” dataset of metastatic breast adenocarcinoma cells (MDA-MB-231) with RGB-markings from a lentiviral gene ontology (LeGO) vector system,^52^ and show that we can effectively use models like Cellpose’s “cyto2”^32^ alongside classical algorithms by applying them independently to each channel and then combining the outputs into a single multilevel contour representation. This approach circumvents the need for extensive retraining or fine-tuning typically required by other approaches.^37, 53^ We combined six segmentation outputs from three image channels — three from Cellpose and three from Otsu^54^ with watershed^55, 56^ — into a single contour and detection map for Ultrack (Fig. 3a). While classical image processing approaches like watershed are generally less precise at splitting cell instances, they provide greater control and allow for the detection of dimmer cells, complementing the more sophisticated Cellpose output. Furthermore, Ultrack can use the three color channels as a feature when associating segments between frames, which helps connect the right segments across time (Fig. 3b).

**Fig. 3.**
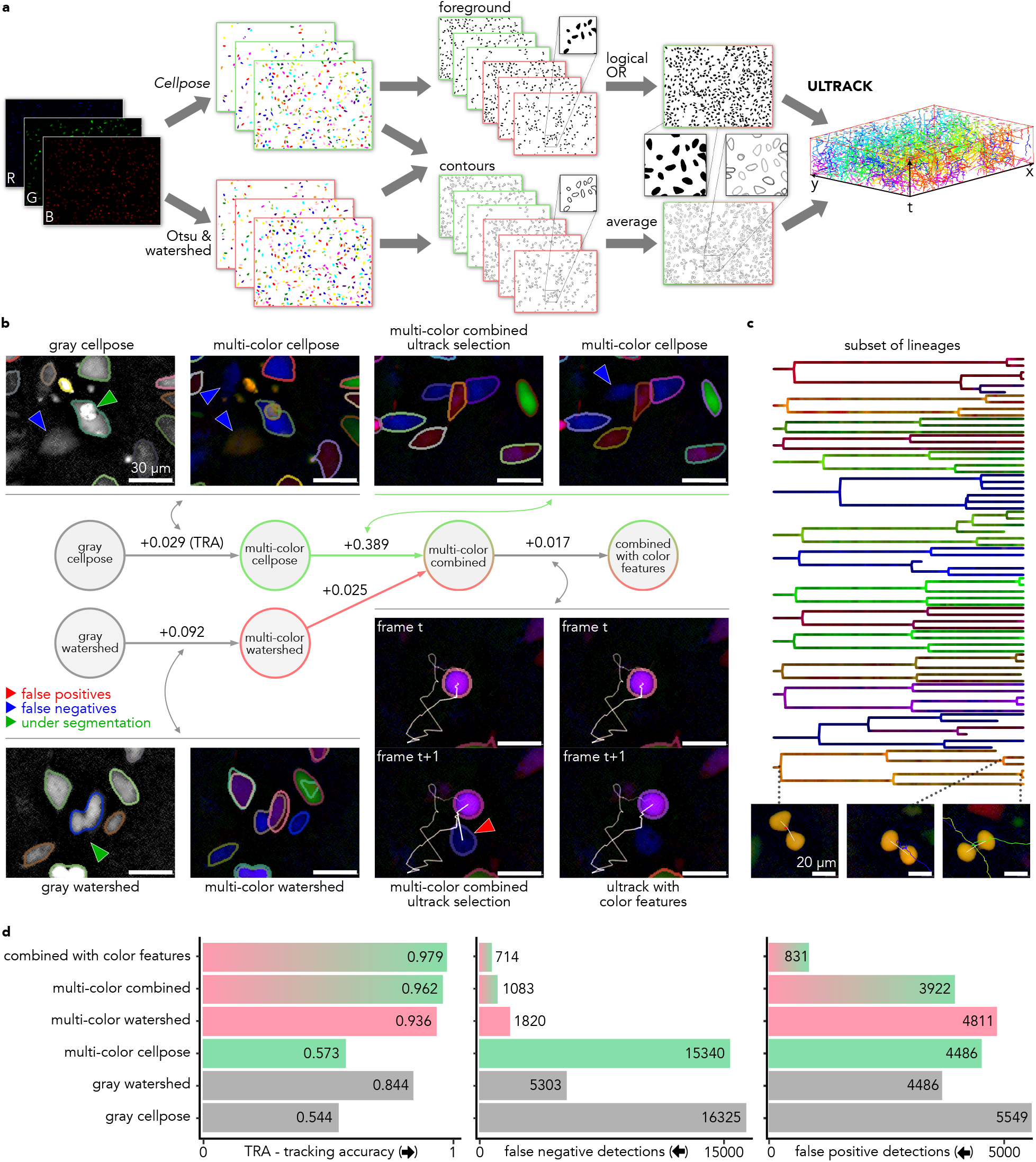
Ultrack enhances multi-color cell tracking by integrating diverse segmentation algorithms. **a**, Multi-channel, multi-algorithm tracking pipeline: Individual labels are generated for each color channel, from which foregrounds and contours are derived and subsequently integrated for joint segmentation and tracking. **b**, Stepwise improvement of segmentation strategies (left to right): (i) individual algorithms applied to grayscale images, (ii) applications to multi-color data, (iii) combination of labels from both algorithms and colors, and (iv) enhancement with color features in association scores. The central diagram illustrates the progression of methods, with numbers indicating improvements in the TRA metric. Colored arrows highlight specific error types. **c**, Representative subset of lineages recovered by Ultrack, showcasing cells’ respective colors and examples of captured division events. **d**, Quantitative evaluation of tracking accuracy for the approaches described in **b**.

We assessed the effectiveness of our multi-channel multi-algorithm approach using progressively more sophisticated strategies: (i) Individual pipelines using Cellpose and watershed algorithms on a grayscale version (maximum intensity across multiple channels); (ii) Each algorithm applied independently on multiple channels; (iii) Combination of outputs from both algorithms in the multi-color configuration (Fig. 3a); (iv) Integration of color features into the association scores to improve linking of segments.

We manually curated 413 cell tracks, creating a goldstandard dataset for benchmarking our results. Demonstrating several key improvements. Incorporating color channels enhances cell distinction compared to grayscale versions, notably reducing under-segmentation (green arrow, Fig.3b, first two columns). The combination of watershed and Cellpose algorithms proves particularly effective, improving the detection of dimmer cells while maintaining Cellpose’s accurate segmentation. Furthermore, color-feature linking improves tracking in ambiguous scenarios (Fig.3b, bottom center). Overall, Ultrack steadily increased accuracy as additional features (*i.e*., segmentation labels, color information) were incorporated. This demonstrates its effectiveness in creating valid lineages (Fig. 3c and Supp. Video 3) from a large pool of segmentation hypotheses without requiring a fine-tuned segmentation model, highlighting its versatility and robustness in multi-color cell tracking scenarios.

### Intensity-based tracking from label-free virtual staining

As demonstrated in the previous section, off-the-shelf segmentation models face challenges when datasets deviate significantly from the training distribution. Users then often resort to time-consuming tasks such as annotating multiple segmentation labels to fine-tune a new model or implementing custom algorithms based on their expertise. This is particularly common because of the high diversity of imaging modalities and datasets typically encountered in Biology. Here, we show that our approach can circumvent these requirements by directly tracking image intensities as if they were multilevel contours, thus eliminating the need for segmentation labels as input.

Our method relies on a single assumption: the intensity map must highlight cell boundaries in some way. This could be achieved through membrane labeling, which delineates cells as a contour map, or through nuclear labeling, which highlights cell interiors. By inverting the latter, we obtain an image where boundaries have higher intensity values than cell interiors.

To evaluate this approach, we tracked A549 cells using quantitative phase imaging (QPI) data acquired with recOrder^59^ on a widefield transmission microscope. Traditional segmentation models like Cellpose perform poorly on this data because QPI was not part of their training set.^57^ We obtained the intensity map using VSCyto2D,^57^ a convolutional neural network for virtual staining (VS) that predicts membrane and nuclear markers from label-free data.

To create the multilevel contour map, we combined both the nuclear and membrane channels by subtracting the normalized and filtered nuclear channel from the filtered membrane channel (Fig. 4a). The foreground was computed by manually thresholding the membrane channel and applying morphological filtering to close holes and remove small objects. We then applied Ultrack to the foreground and the intensity map encoding cell contours (see Supp. Video 4).

**Fig. 4.**
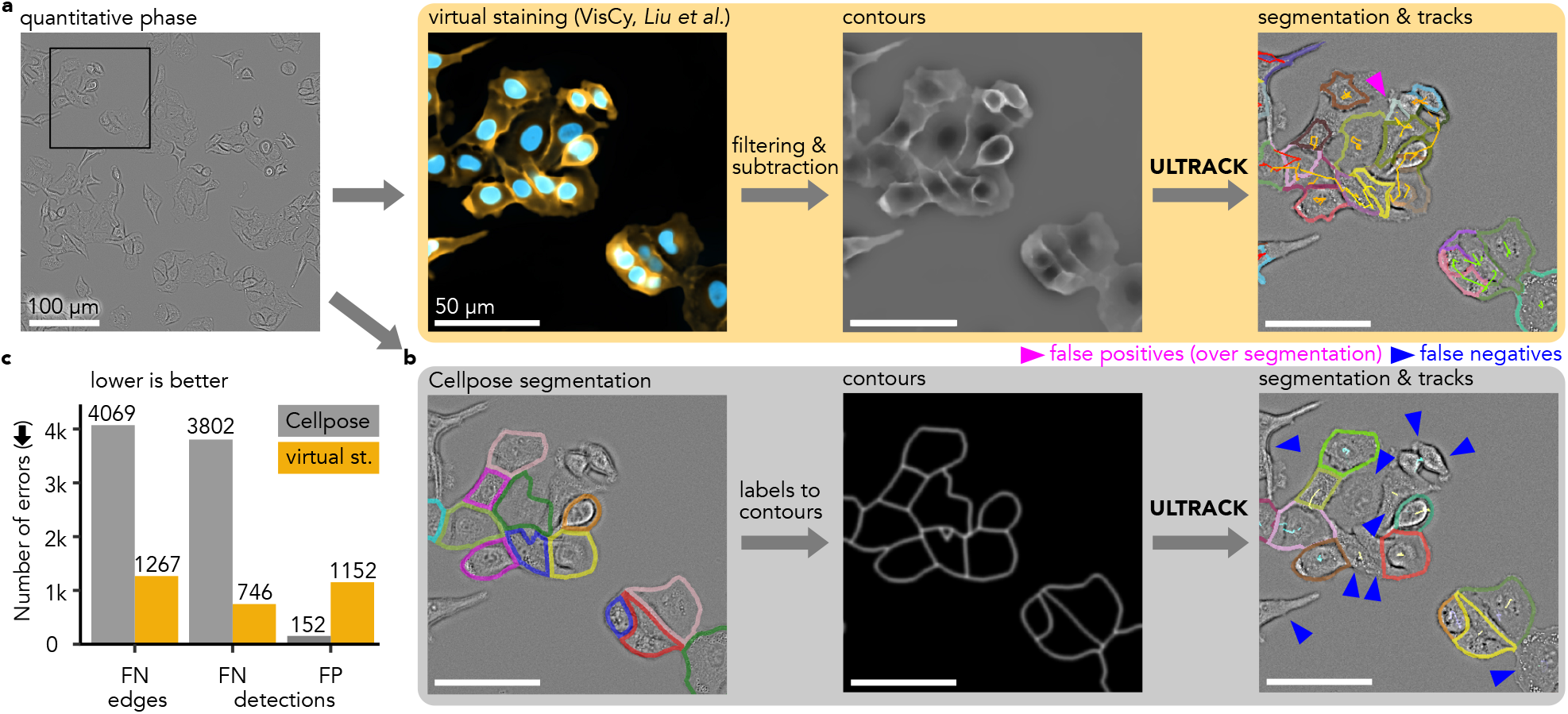
Ultrack improves cell tracking in label-free imaging by leveraging virtual staining. **a**, Pipeline for intensity-based joint segmentation and tracking from label-free quantitative phase imaging: Quantitative phase image (top left) is processed by VisCy^57^ to generate virtual staining of nuclei and membrane (second from left). These virtual stains are used to derive contours (third from left) and foreground masks (not shown) through filtering, subtraction, and thresholding. Ultrack uses these inputs to perform tracking (right). **b**, We compare the above against first applying a Cellpose segmentation to the Quantitative phase image (bottom left), then deriving contours (and foreground mask, not shown), and then applying Ultrack (right). False positives (over-segmentation) are indicated in magenta, and false negatives in blue. **c**, Quantitative comparison of tracking errors using Cellpose^58^ segmentations (gray) versus virtually stained membranes (orange) as input. The bar chart shows the number of false negative (FN) edges, false negative (FN) detections, and false positive (FP) detections for each method.

For benchmarking, we tracked the virtual stained nuclear channel and manually curated a set of 247 tracklets of lineages starting in the first frame to the last frame. The cell centroids of these tracks were used to create reference gold-standard datasets following the Cell Tracking Challenge (CTC) format.

We compared against Cellpose with “livecell cp3” weights^58^ applied to the QPI channel, followed by tracking. With Ultrack applied to the combined membrane and nuclear VS channels, we detected most of the cells, Fig. 4b-c, while the off-the-shelf model failed to recognize them. We missed 2802 fewer associations between cells (*i.e*. FN edges) and 3014 cell detections (*i.e*. FN detections) than VS with Ultrack, at the cost of detection of 1000 false-positive cells.

This shows that tracking from image intensity directly, in this case from virtual staining, offers a competitive alternative to supervised segmentation and tracking methods, which often fail when presented with unseen imaging modalities, such as label-free quantitative polarization, Zernike phase contrast, DIC, or for different cell types. Thus eliminating the need for extensive human annota-tion to train segmentation models.

### Improving tracking by temporal registration

Imaging systems must balance multiple factors, including the field of view size, spatial resolution, imaging speed, and depth.^60^ A critical limitation often encountered is acquisition speed (*i.e*., temporal resolution), which hampers cell tracking performance. This is especially problematic when cells move rapidly and have similar appearances, complicating the assignment of cells between consecutive frames.

Registration of adjacent frames often compensates for limited time resolution. However, collective cell motion within growing or deforming tissues frequently demonstrates local coherence that does not conform to simple linear or affine transformations. This is due to the complex dynamics of some biological processes. In such cases, non-linear registration, such as movement vector fields,^61, 62^ is more appropriate. This approach effectively minimizes apparent motion when cells present distinct migration patterns.

Computing non-linear registration can be extremely time-consuming or require complex, specialized algorithms. To address this, Ultrack provides its own graphicshardware-accelerated (GPU) routines to compute vector fields for 2D and 3D time-lapses. These routines are based on automatic differentiation from deep-learning models,^63^ offering efficient computation without sacrificing accuracy (see Methods for details). The movement vector fields can be integrated into the tracking step, applying local movement to each candidate segmentation. This approach avoids altering the original image data. Moreover, Ultrack accepts any vector fields represented as a grid, allowing users to utilize their preferred registration methods.

We evaluated the effectiveness of using vector fields with Ultrack on the Tribolium Castaneum cartography dataset^64^ from the Cell Tracking Challenge (CTC). This dataset features nuclear-labeled embryo cells imaged using selective plane illumination microscopy (SPIM). The epithelial cells’ ellipsoidal surface was transformed into planar cartography using the ImSAnE toolbox^65^ to facilitate downstream analysis. However, this cartographic transformation introduced distortion and excessive motion on the polar boundaries, as shown in Fig. 5a and Supp. Video 5.

**Fig. 5.**
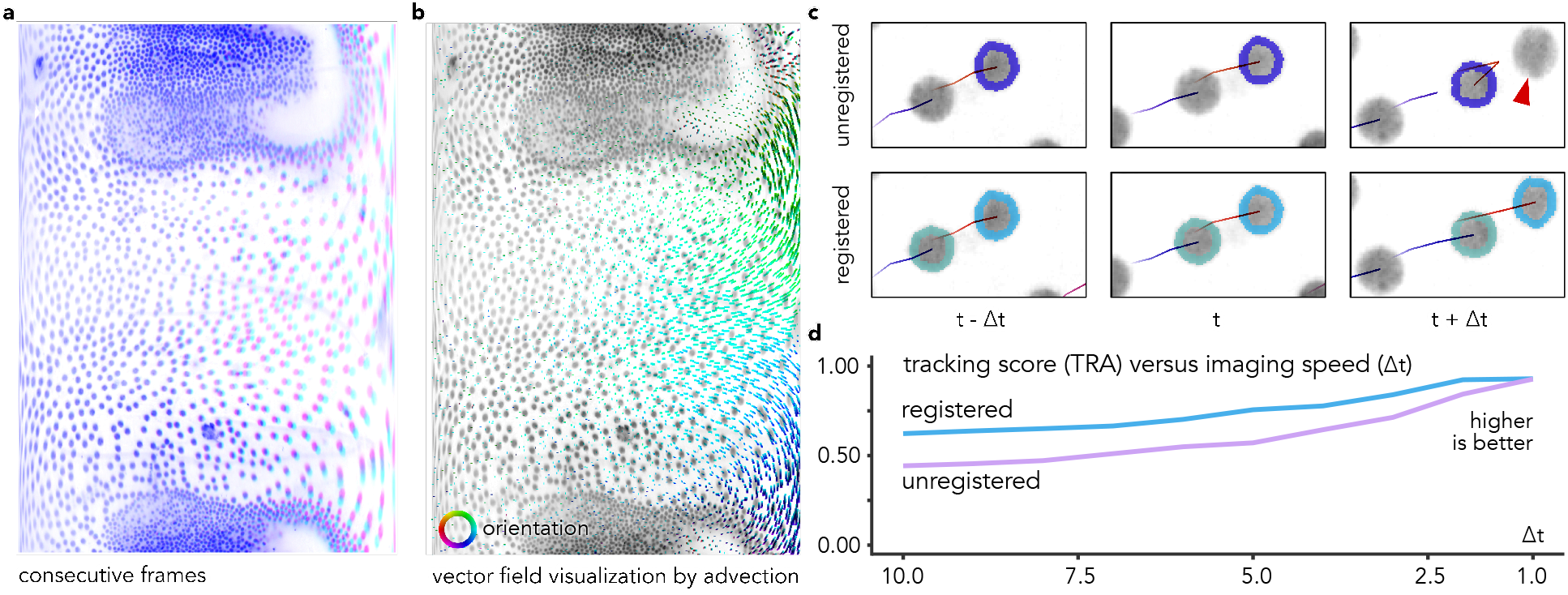
Enhancing tracking accuracy through temporal registration. **a**, Consecutive frames, *t* (magenta) and *t* + 1 (cyan), highlighting divergent motion along the right boundary. **b**, Vector field generated by non-linear registration, colored by orientation. **c**, Qualitative comparison of cell identity preservation: top row shows a cell identity switch in unregistered frames, identity switch indicated by red arrow, bottom row demonstrates improved consistency with flow registration. **d**, Comparative analysis of tracking accuracy for registered and unregistered data across different cell velocities (Δ*t*), illustrating enhanced performance with registration.

We compared the tracking accuracy with and without flow registration (Fig. 5b) at various temporal resolutions, starting by skipping ten frames (Δ*t* = 10) up to the original framerate (Δ*t* = 1), as illustrated in Fig. 5c-d. The results show: (i) At Δ*t* = 10 tracking accuracy improved from 0.443 to 0.623 with flow registration; (ii) At Δ*t* = 5 accuracy increased from 0.571 to 0.756; (iii) At original time resolution (Δ*t* = 1) similar results were obtained (0.927 vs. 0.929). Fig. 5c provides a specific example where the flow field achieves the correct result, while a cell identity switch occurs in the unregistered case.

These results demonstrate that Ultrack’s temporal registration capability enhances tracking accuracy, particularly when dealing with complex cell movements or lower temporal resolution datasets.

All these features — including its ability to handle multiple segmentation hypotheses, integrate multi-channel data, leverage intensity-based tracking, and apply temporal registration—collectively enable robust cell tracking in complex biological contexts. These capabilities are particularly valuable when confronting the challenges posed by dense, three-dimensional tissues with dynamic cellular behaviors. To demonstrate the power and versatility of this approach, we next evaluated Ultrack’s performance on one of the most demanding applications in bioimage analysis: tracking cells in developing embryos.

### Ultrack achieves state-of-the-art cell tracking in developing embryos

The Cell Tracking Challenge (CTC) is a long-standing benchmark for cellular segmentation and tracking methods.^11^ It provides participants with training datasets to tune and train their algorithm parameters; the test sets are concealed and only accessible to the organizers. This structure prevents information leakage between the training and testing phases and offers an unbiased evaluation of each algorithm’s performance. We assessed our method using their three 3D whole-embryo datasets: the *C. elegans* (worm),^66^ *D. melanogaster* (fly),^31^ and *T. castaneum* (beetle) embryos.^64^

The worm dataset includes manually curated lineage data and cell coordinates, as well as silver-standard segmentation annotations derived from a consensus among prior challenge submissions.^67^ Using these annotations, we trained a U-Net model^45, 68^ that predicts two outputs per voxel: (i) whether the voxel is on a cell contour and (ii) whether that voxel is in the foreground. We optimized these predictions using three dice losses:^69^ (i) binary foreground prediction, (ii) binary cells’ contour prediction, and (iii) U-Net’s intermediate cell contour predictions.^70^ The cells’ contours and foreground were then used as inputs for Ultrack. As shown in Table 1, Ultrack achieves a top combined segmentation and tracking score of 0.844.

**Table 1.** Segmentation and tracking scores for 3D embryo datasets of the Cell Tracking Challenge. Ultrack results, as of August 2024, are highlighted in bold. Ultrack obtains the highest combined segmentation and tracking score (CTB) for all three datasets and is the method with the best balance between segmentation (SEG) and tracking (TRA) metrics.

| Dataset | Rank | combined score<br>(CTB) | tracking score<br>(TRA) | segmentation score<br>(SEG) |
| --- | --- | --- | --- | --- |
| Worm<br>*C. elegans* | 1<sup>st</sup> | **0.844**<sup>ultrack</sup> | 0.987<sup>MPI</sup> | 0.759<sup>MU</sup> |
|  | 2<sup>nd</sup> | 0.829<sup>KTH</sup> | 0.979<sup>JAN</sup> | 0.729<sup>KIT</sup> |
|  | 3<sup>rd</sup> | 0.808<sup>KIT</sup> | 0.975<sup>IGFL</sup> | 0.722<sup>KTH</sup> |
|  | 4<sup>th</sup> | - | **0.967**<sup>ultrack</sup> | **0.722**<sup>ultrack</sup> |
| Fly<br>*D. melanogaster* | 1<sup>st</sup> | **0.708**<sup>ultrack</sup> | **0.802**<sup>ultrack</sup> | **0.613**<sup>ultrack</sup> |
|  | 2<sup>nd</sup> | 0.617<sup>KTH</sup> | 0.785<sup>JAN</sup> | 0.613<sup>KTH</sup> |
|  | 3<sup>rd</sup> | 0.591<sup>JAN</sup> | 0.785<sup>MPI</sup> | 0.397<sup>JAN</sup> |
| Beetle<br>*T. castaneum* | 1<sup>st</sup> | **0.841**<sup>ultrack</sup> | 0.955<sup>MPI</sup> | **0.746**<sup>ultrack</sup> |
|  | 2<sup>nd</sup> | 0.804<sup>MPI</sup> | **0.936**<sup>ultrack</sup> | 0.684<sup>RWTH</sup> |
|  | 3<sup>rd</sup> | 0.785<sup>RWTH</sup> | 0.886<sup>RWTH</sup> | 0.654<sup>MPI</sup> |

In general, training data for segmentation is not always available nor easily obtained, which is a typical situation users face. This is the case for the fly and beetle datasets, which lack ground truth to train segmentation models. Instead, we asked whether we could forgo machine learning and use classical image-processing techniques for segmentation. We opted for a straightforward method. First, we detected the foreground using a Difference of Gaussians filter and applied Otsu’s threshold^54^ to select the foreground voxels. Next, we used image intensity as a proxy for cell boundaries by inverting and normalizing these intensities between 0 and 1, so dimmer voxels, which are more likely to be the cells’ contours, represent the highest values.

Combining this simple image-processing approach with Ultrack’s handling of multiple hypotheses implicit in the contour representation yields noteworthy results compared to competing approaches that require training data. As shown in Table 1, for the fly dataset, Ultrack achieved a combined segmentation and tracking score of 0.708, compared to 0.617 for the next best method. Similarly, for the beetle dataset, Ultrack scored 0.841, while the next best result was 0.804 (see Methods for additional details). These outcomes demonstrate Ultrack’s capability to perform well even when deep-learning models are unavailable. See Supp. Video 6 for recordings of the results.

### High-quality cell tracking ground-truth via sparse fluorescence labeling

While the Cell Tracking Challenge (CTC) provides a valuable unbiased benchmark for evaluating segmentation and tracking, it relies primarily on human-generated annotations. These annotations, while crucial, are inherently limited in scale and potentially in quality due to the challenge of manually annotating the latest-generation 3D live microscopy datasets.^1, 2, 6, 7, 9, 15, 16^ Such datasets often span hundreds of gigabytes to terabytes, recording thousands of cells over hundreds or more time points, making manual annotation a task that is not only time-consuming but often beyond human capability. Moreover, in densely labeled or multi-dimensional datasets, distinguishing individual cells can be highly uncertain and sometimes impossible.^15, 16, 24, 31, 64^

Encouraged by Ultrack’s performance on the CTC, we developed a higher-fidelity ground truth dataset that could overcome these limitations and potentially provide a more challenging benchmark of a crowded and dynamic cellular environment from a developing zebrafish embryo. We applied a dual labeling protocol that combines ubiquitous fluorescence nuclear labeling (H2B-mNeonGreen) with sparse random nuclear labeling at a distinct wavelength (pMTB-ef1-H2B-mCherry). Sparse labeling was accomplished by early-stage microinjection of a marker (Tol2 transposase-mediated integration, see Methods for details) that propagates to daughter cells upon division, effectively recapitulating their lineage (Fig. 6a).

**Fig. 6.**
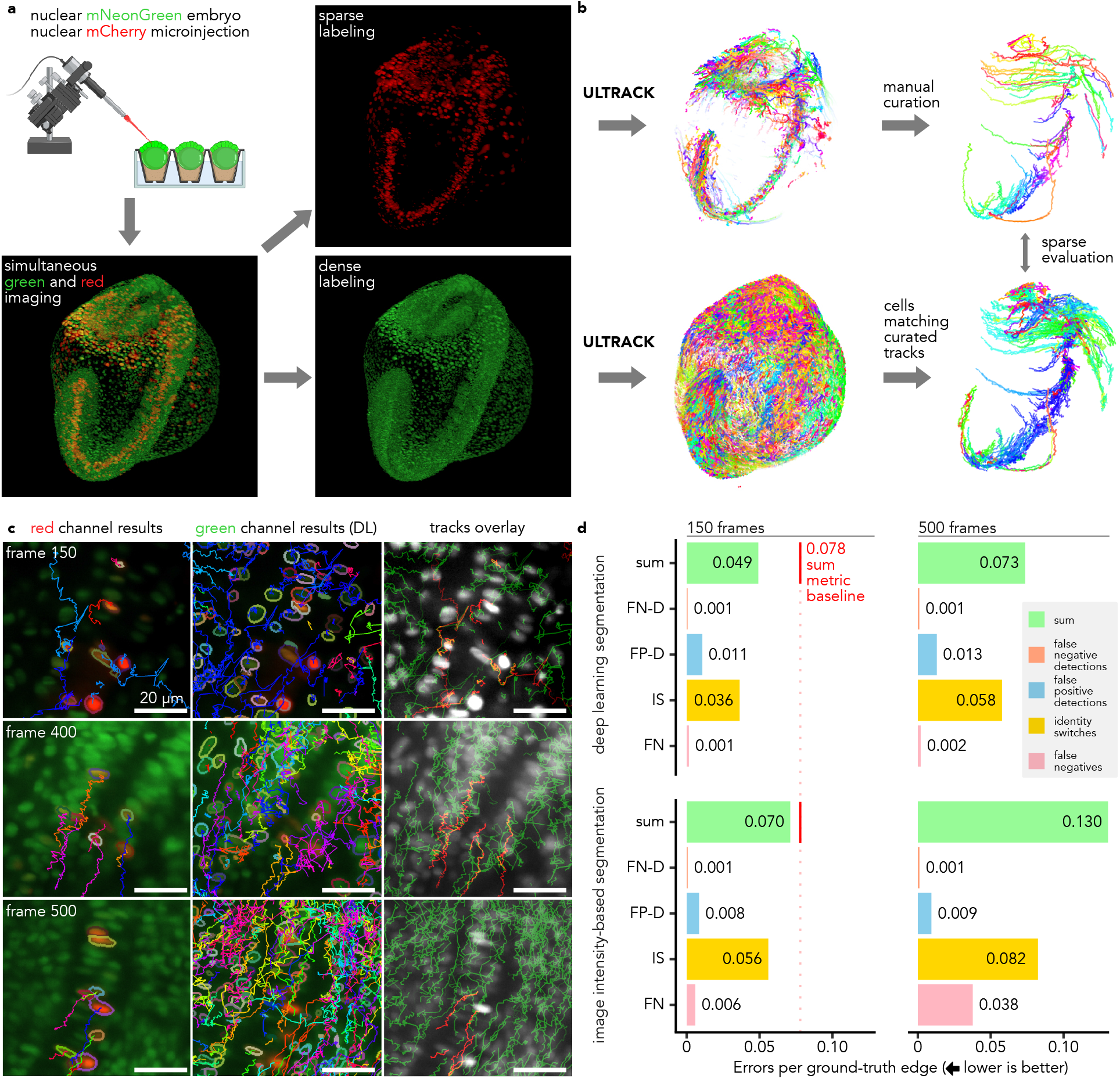
Sparse fluorescence labeling enables high-fidelity tracking validation over extended time-lapses. **a**, Validation pipeline using sparse fluorescence labeling: (i) Embryos with ubiquitous green nuclear labeling (H2B-mNeonGreen) are injected with DNA plasmids expressing red fluorescent protein pMTB-ef1-H2B-mCherry; a single embryo is then selected for imaging. (ii) Simultaneous imaging of red and green channels. **b** (iii) Independent tracking of cell nuclei in both channels. (iv) Selection and manual curation of a subset of cell nuclei tracks from the red channel. (v) Comparison of curated tracks to those obtained from ubiquitous labeling. **c**, Comparative visualization of red (first column) and green (second column) channel segmentation and tracking results at different time points, and combined results overlaid grayscale intensity colored in red and green (last column); cell density and tracking difficulty increase over time (top to bottom), **d**, Quantitative evaluation of tracking accuracy under different conditions and durations: using deep learning for cell contour estimation (first row) vs. direct image-intensity-based segmentation (second row); measurements taken up to the first 150 frames (first column) and 500 frames (second column). Metrics shown are proportions of false-negative edges (FN), identity switches (IS), false-positive divisions (FP-D), false-negative divisions (FN-D), and their sum. The red vertical line indicates the baseline sum error rate for 150 frames of 0.078 for comparison with Malin *et al*.^24^

The sparsity of this second channel significantly simplifies tracking and allows for rapid curation into a set of high-quality, or “platinum-level,” annotations (Fig. 6b). Cells in the sparse channel were automatically tracked using Ultrack and then manually curated to ensure the highest quality standards (Fig. 6b–top). This curation process involved selecting long, green-overlapping, high-quality lineages, conducting visual inspections, and making necessary corrections, resulting in 152 annotated tracklets spanning from 85 to 521 frames. This methodology not only simplifies the manual curation process but also reveals cell trajectories that are challenging to annotate manually using conventional methods on dense datasets, see Supp. Video 7.

As evident in Fig. 6c, tracking is easier at earlier stages of development when cells are less numerous, less densely packed, and when the optical properties of the tissue are more favorable for imaging. Despite these challenges, cell nuclei in the sparse channel are consistently easier to track compared to the dense channel (Fig. 6c, first row).

We used these high-quality sparse-channel-derived tracking labels to evaluate Ultrack’s accuracy on the ubiquitously labeled channel. We used two segmentation approaches as in:^45^ (i) image processing to produce foreground masks and nuclei boundaries directly and (ii) a convolutional neural network to generate foreground and contour images. Hence, the Ultrack setup was kept the same, and only the input foreground and contour maps were modified. Figure 6c compares the tracking results using the sparse channel (red) and the ubiquitous channel (green) using the deep-learning-based contour map, approach (ii), for three frames (150, 400, and 500), showcasing the disproportionate difficulty of tracking cells in the dense versus sparse channel.

To quantitatively assess the performance, we evaluated our automatic dense tracking results against the manually curated sparse tracking over two time periods, Fig 6d: the first 150 frames as the previous work^24^ and an extended period of 500 frames, nearly covering the entire timelapse. Assessing tracking performance with five metrics,^24^ all expressed as proportions relative to the total number of annotated edges: (i) FN: false-negative edges (*i.e*. missing connections between cells), (ii) IS: identity switches (*i.e*. incorrect association between cells), (iii) FP-D: falsepositive divisions, (iv) FN-D: false-negative divisions, and (v) the sum error rate that combines the first four metrics.

As expected, tracking accuracy drops for longer imaging periods (500 versus 150 frames, see Fig. 6d), reflecting the growing complexity of the developing organism already apparent in Fig. 6c. Notably, Ultrack with a deeplearning-based input, which benefits from available training data, consistently outperforms plain image processing contour and foreground maps, maintaining comparable accuracy over 500 frames to what the image intensity-based approach achieves in just 150 frames. This performance demonstrates the robustness of the deep-learning method in maintaining high tracking accuracy over extended periods of embryonic development, even as the tracking task becomes increasingly challenging.

To contextualize our results within recent advances in cell tracking for embryo-scale 3D datasets, we compared our findings to those reported by Malin *et al*.^24^ Their study, which used a similar multi-view light-sheet microscope setup,^71^ established a baseline sum error rate of 0.078 for tracking neuronal cells in zebrafish embryos during 150 frames. As shown in Fig. 6d, Ultrack achieves a lower proportion of sum errors for both deep learning and intensity-image-based contour maps on 150 frames (0.049 and 0.070, respectively). Moreover, when using deep learning, we maintain accuracy (0.073) comparable to our baseline (0.078) even over a much more challenging 500-frame period. This extended performance is particularly noteworthy, as it underscores the robustness of our method in maintaining high tracking accuracy over prolonged periods of embryonic development despite increasing tissue complexity, cell density, and imaging artifacts. These results not only validate the effectiveness of our dual-channel approach but also highlight Ultrack’s potential to push the boundaries of long-term cell tracking in complex biological systems.

### Ultrack scales to terabyte-scale high-resolution largefield-of-view light-sheet datasets

To demonstrate Ultrack’s scalability and performance on massive datasets, we employed it to recover cell lineages of nuclear-labeled zebrafish embryos^9^ acquired using the fast (1 stack per minute), high-resolution (20x magnifica-tion and 1.0 NA Olympus objective), and large-imagingvolume (≥ 1 mm^3^) DaXi light-sheet microscope,^6^ producing uncompressed time-lapse data ranging from 1.7 to 3.7 terabytes, capturing 8.6 to 13.2 hours of zebrafish development (Fig. 7a,b).

**Fig. 7.**
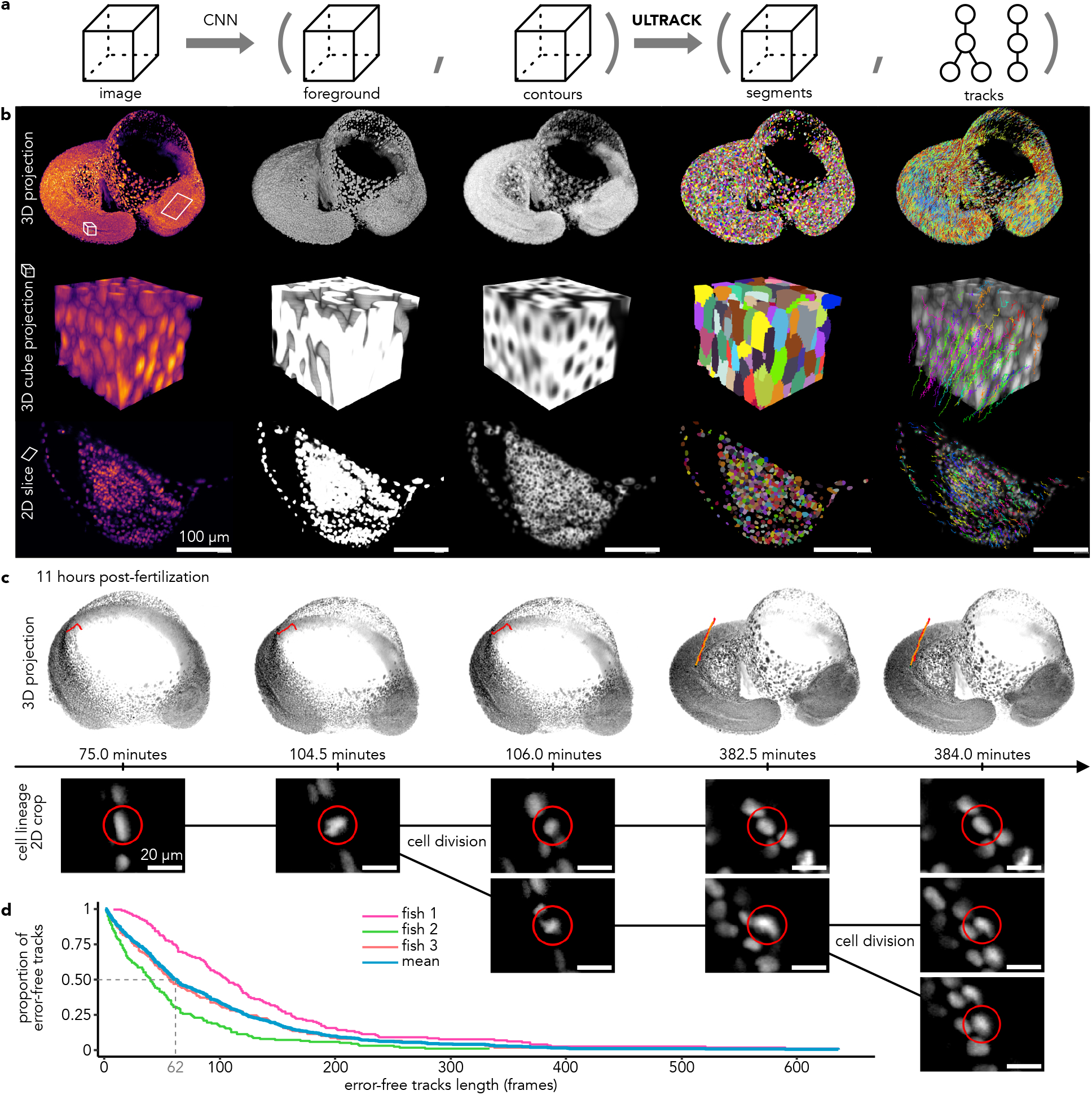
Multi-terabyte cell tracking of zebrafish embryo. **a**, Pipeline for joint segmentation and tracking using a convolutional neural network and Ultrack. **b**, Intermediate results at each stage: original image (first column); predicted foreground (second column) and contour probabilities (third column); Ultrack segmentation (fourth column) and tracking results (fifth column). **c**, Detailed views of fully automated lineage reconstructions, showcasing two rounds of cell divisions and their corresponding 2D slices. **d**, Distribution of the proportion of error-free tracks for varying lengths across three embryos. Indicating that, on average, 50% of the lineages are error-free for at least 62 frames.

Our workflow began with training a U-Net to predict nuclei pixels (foreground) and their contours (Fig. 7a,b). We then applied Ultrack to these network predictions. Leveraging Ultrack’s command-line interface (CLI) and SLURM,^72^ we processed these massive datasets within hours using distributed computing in a high-performance computing (HPC) environment. For example, a 3.7 TB dataset (791 × 448 × 2174 × 2423 voxels, T × Z × Y × X) was processed in approximately 8.2 hours (Fig. 1e).

Ultrack’s computational scaling extends across computing environments, from high-performance clusters to offthe-shelf laptops. Ultrack supports out-of-core processing: it is able to process arbitrarily large datasets, even those significantly larger than available memory (RAM) — as long as a single image (2D or 3D) can fit into memory (see Methods for details). We could process the same 3.7 TB dataset on a 64 GB RAM laptop in 5 days, accessing data remotely via a 1 Gbps connection (Fig. 1e).

The tracking resulted in millions of segments across hundreds of time frames. To assess performance, we manually evaluated time-lapses until the first tracking mistake, recording the duration of error-free tracks. We employed a stratified sampling strategy, manually annotating distinct regions (head, body, tail bud, and presomitic mesoderm) at various time points and image qualities. From these strata, we randomly sampled approximately 140 lineages per embryo (n = 3). Figure 7d shows each fish’s cumulative proportion of error-free tracks and their mean. On average, it took 62 frames for a tracking error to appear in 50% of the lineages, meaning that for a period of approximately 1 hour of development (60 frames), half of the tracks are completely error-free, demonstrating Ultrack’s robust performance over extended periods. Figure 7c and Supp. Video 8 provides a detailed view of an automatically reconstructed lineage, capturing two rounds of cell divisions and illustrating Ultrack’s ability to maintain accurate tracking through multiple cell generations.

These results underscore Ultrack’s capacity to efficiently process and accurately track cells in multi-terabyte, high-resolution datasets utilizing HPC resources or standard hardware. This scalability and versatility make Ultrack a powerful tool for analyzing large-scale developmental processes at unprecedented resolution and scale.

### Ultrack attains near-perfect cell tracking of organ-scale 3D datasets

Cell tracking is often crucial in scenarios with fewer cells, where perfect lineage reconstruction is necessary to uncover precise details of cell behavior, such as division, migration, and death.^73, 74^ To evaluate Ultrack’s performance in such a context, we chose the zebrafish neuromast as our model system, given its well-defined structure and importance in studying sensory organ development. Previous efforts in segmenting and tracking cells in this organ included manual tracking of cells^75^ or semi-automated segmentation of single frames.^76^ To our knowledge, no other study has reported the fully automatic segmentation and tracking of cells in this complex 3D organ for extended time-periods.

We imaged a zebrafish neuromast for 42 hours using membrane and nuclei markers. Imaging was performed on an Olympus IX83 microscope equipped with a microlens-based, super-resolution spinning disk confocal system (VT iSIM, VisiTech International), using a 60x 1.3 NA silicone-oil objective. This setup yielded a dataset of 500 frames in two colors, with volumes of 73 × 1024 × 1024 voxels, at a resolution of 250 × 76.6 × 76.6 µm per voxel. Frames were captured every 5 minutes.

To ensure optimal segmentation accuracy, we finetuned a 3D Cellpose model^32^ to segment cells using both color channels. We then converted the Cellpose predictions into foreground and contour maps for input into the Ultrack algorithm (Fig. 8a). This approach allowed us to reconstruct lineages spanning the entire time-lapse, capturing diverse cellular events such as migration within the organ, cell division, and cell death (Fig. 8b-c and Supp. Video 9).

**Fig. 8.**
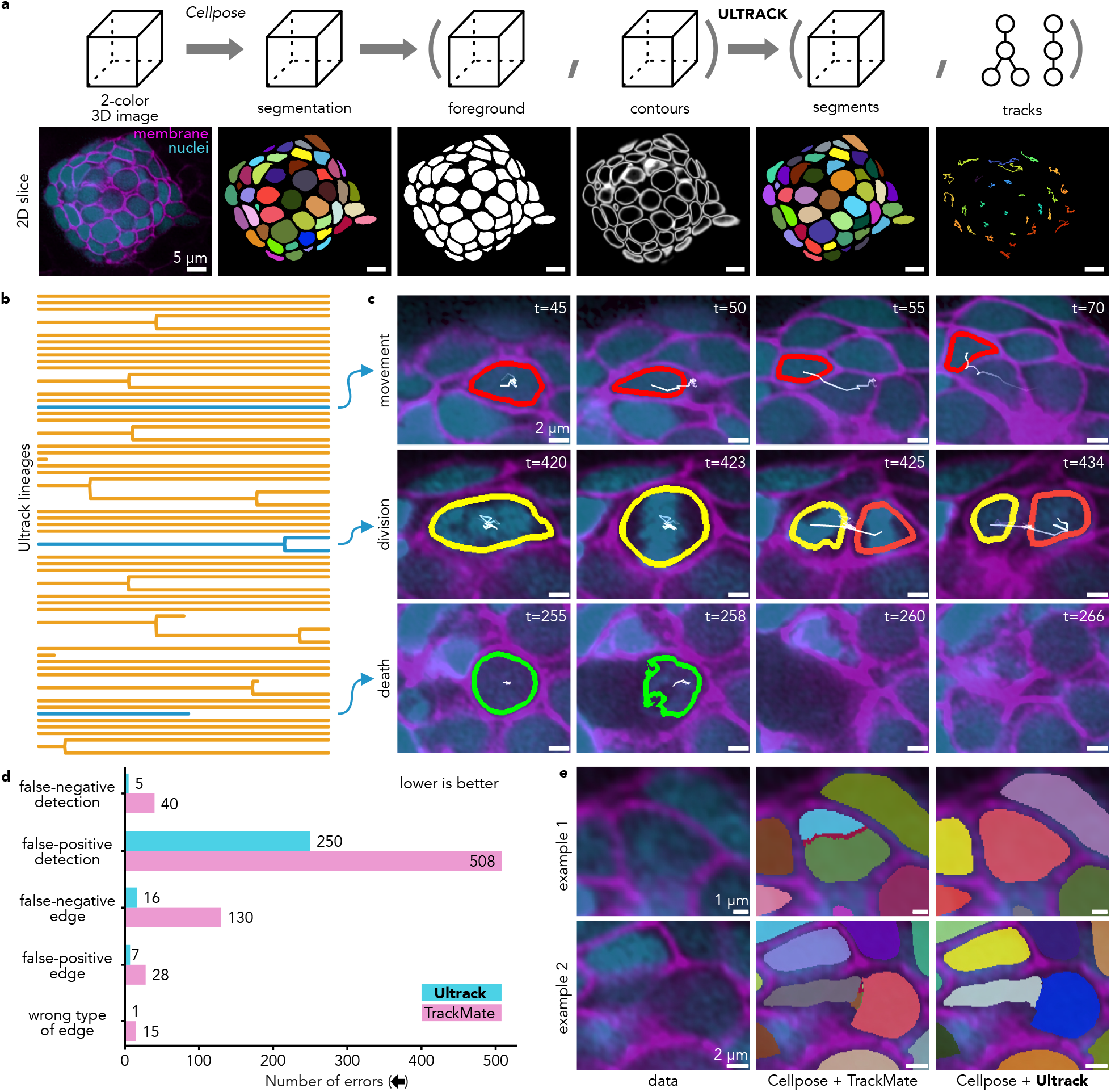
Near-perfect nuclear- and membrane-based 3D tracking of zebrafish neuromast cells. **a**, Pipeline for joint segmentation and tracking using Cellpose and Ultrack, showing 3D schematics (top) and 2D slice examples (bottom) at each stage: original 2. color image, Cellpose segmentation, foreground and contour probabilities, Ultrack segmentation, and tracking results. **b**, Dendrogram recovered by Ultrack for 71 cells over 500 frames (41.7 hours), with orange lines representing cell lineages and blue highlights indicating examples detailed next: **c**, Time-lapse images illustrating cell movement (red outlines), division (yellow/orange outlines), and death (green outlines). **d**, Quantitative comparison between Ultrack (blue) and TrackMate (pink). The bar chart shows error counts for various metrics, with Ultrack obtaining 442 fewer mistakes than TrackMate. **e**, Two examples demonstrating Ultrack’s robustness to over-segmentation, showing the original image (left), fragmented TrackMate results (middle), and accurate Ultrack segmentation and tracking (right) despite imperfect Cellpose input.

To assess Ultrack’s accuracy, we visually inspected the generated lineages and manually corrected any errors, creating ground-truth reference lineages. Using the Cell Tracking Challenge (CTC) evaluation routines, we quantified Ultrack’s performance (Fig. 8d). Ultrack achieved a TRA (tracking accuracy) score of 0.9989, demonstrating its high precision in this challenging scenario.

We compared Ultrack’s performance against TrackMate,^37^ using identical Cellpose segmentations as input. To ensure a fair comparison, we first optimized both Ultrack’s and TrackMate’s parameters using a different 50-frame dataset. Key parameters tuned included minimum and maximum cell size and the maximum movement. We then applied these optimized settings to the full 500-frame dataset reported here.

In this case, Ultrack’s tracking results exhibit a more than 2-fold reduction in errors (279 vs. 721), with 442 fewer mistakes needing manual correction when compared to TrackMate. The primary source of errors in TrackMate’s results stemmed from imperfect Cellpose segmentations (Fig. 8d). Despite TrackMate’s segmentation filtering options, it struggled to correct issues such as oversegmentation and false merges. In contrast, Ultrack’s approach naturally incorporates these imperfect segments into its hypotheses, allowing for more accurate merging and tracking (Fig. 8e).

These results demonstrate Ultrack’s ability to maintain high tracking accuracy in this complex, long-term imaging scenario and, thus, its utility for detailed studies of cellular behavior and tissue dynamics.

### Democratizing high-performance cell tracking at all skill levels

Cell tracking software often represents one of the most complex components in image analysis pipelines, primarily due to the necessity of handling the time dimension— a factor frequently overlooked in segmentation and other image processing operations. Moreover, it operates on non-matrix data (*i.e*. track lineages), where efficient data processing routines are less prevalent. This complexity has historically restricted accessibility to a broad user base, with only a few longstanding solutions like TrackMate^37^ and others,^21, 22, 31^ alongside specialized solutions for handling larger datasets.^24^

Ultrack addresses these challenges by offering a versatile, user-friendly solution that caters to a wide range of users, from biologists with minimal programming experience to machine learning experts requiring customizable inputs. By providing multiple interfaces and scalable processing capabilities, Ultrack democratizes access to advanced cell tracking technology, enabling efficient analysis of datasets of various sizes across different computational resources (see Fig. 1e).

At the heart of Ultrack’s accessibility is its multiinterface approach. Ultrack offers two options for users who prefer a graphical interface: napari and Fiji. The *napari* plugin (Fig. 9a) integrates transparently with the Python ecosystem, providing a comprehensive set of features for non-coding users. This interface offers graphicshardware-accelerated pre-processing, vector-field flow computation, and full configuration of Ultrack parameters. Complementing this, the Fiji plugin^35^ (Fig. 9b) exposes Ultrack’s algorithm to Fiji’s mature ecosystem and broad user base, allowing seamless integration with existing Fiji workflows and ensuring compatibility with popular tools like TrackMate^37^ and Mastodon.^38^ Supp. Video 10 presents them in action.

**Fig. 9.**
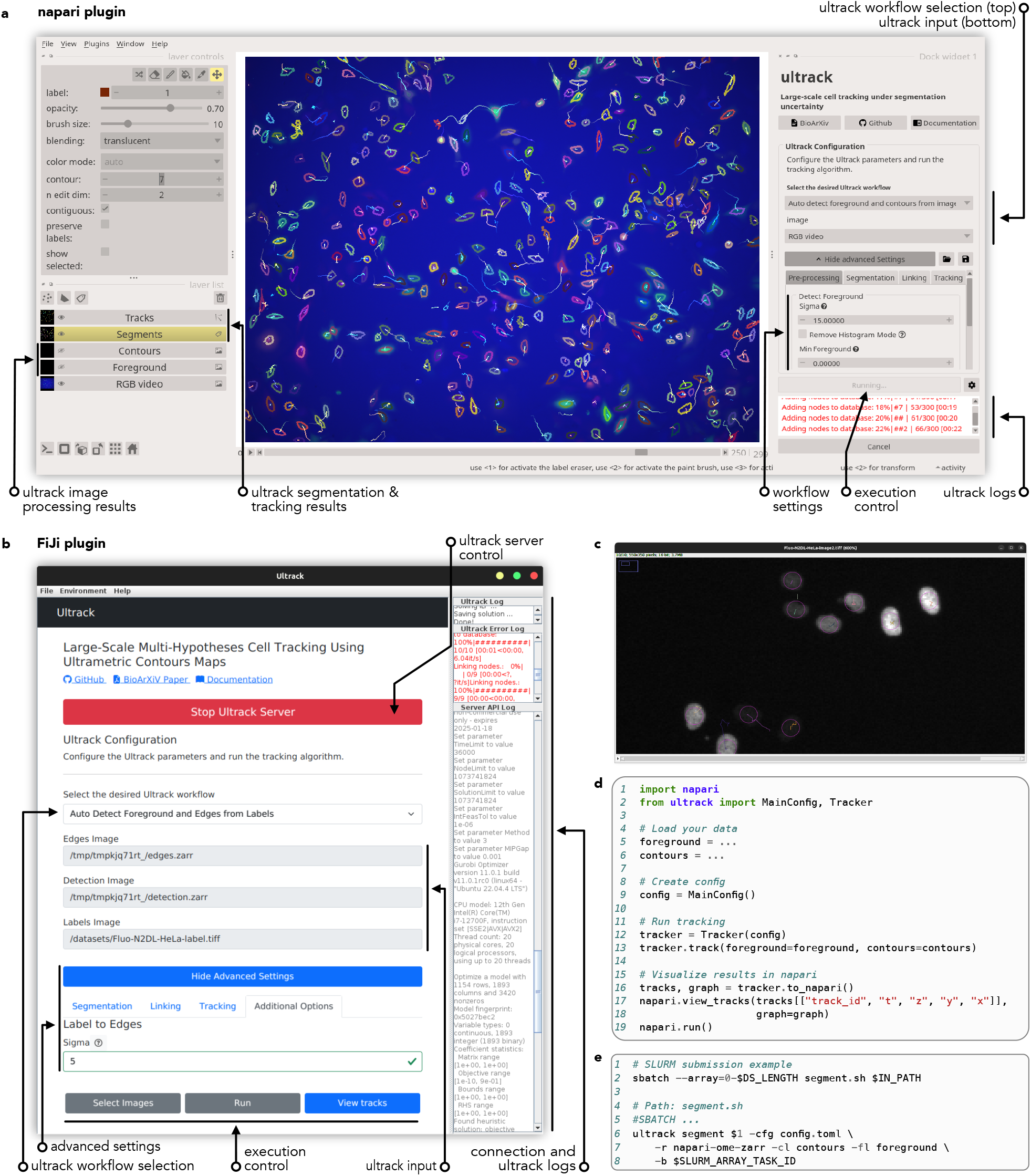
Ultrack’s multi-interface approach enables cell tracking for diverse user needs and computational environments. **a**, Napari plugin interface showcasing multi-color automatic cell tracking results. The interface includes workflow selection, input configuration, image processing results, segmentation and tracking visualization, workflow settings, and execution control. **b**, Fiji plugin interface guiding users through workflow selection, input specification, and advanced settings. It also provides execution control and displays Ultrack server logs. **c**, Example of image data overlaid with cell detections and tracks as viewed in the Fiji interface. **d**, A concise Python API example (18 lines) demonstrating data loading, configuration, tracking execution, and result visualization in *napari*. **e**, Sample SLURM submission script for Ultrack’s CLI, illustrating its integration with high-performance computing environments for distributed processing.

For users comfortable with programming, Ultrack provides a user-friendly Python API (Fig. 9c,d) that exposes all of its functionalities, allowing for easy customization and reproducible workflows. This API accepts various array formats (numpy, zarr, dask, etc.), and can export data into multiple file formats, including networkx,^77^ tables (*i.e*., data frames),^78^ Cell Tracking Challenge,^11^ and TrackMate^37^ formats, enhancing its interoperability with other tools and workflows. Additionally, Ultrack offers a command line interface (CLI, see Fig. 9e) designed for batch processing and streamlined workflows, which supports distributed processing in high-performance computing environments. This CLI accommodates any file format supported by existing *napari* reader plugins, eliminating the need for data conversion — a crucial feature for handling large datasets.

## DISCUSSION

Ultrack advances cell tracking by offering a robust, scalable solution for analyzing complex cellular dynamics across various scales and dimensions. Its integration of segmentation and tracking, leveraging multilevel contour maps (UCMs), enables effective analysis in challenging scenarios where previous methods often struggled.

The method’s versatility is demonstrated through its top-scoring performance in 3D embryonic cell tracking challenges, even when using non-learned classical image processing algorithms. This ability to perform well without training data or fine-tuning addresses a critical challenge in the microscopy field, where 3D annotations are often scarce.^11^ Furthermore, Ultrack’s capacity to combine results from parameter sweeps eliminates the common problem of identifying optimal segmentation parameters. This flexibility extends to working directly with image intensities thus circumventing the need for explicit segmentation in certain scenarios, for example, virtual staining of label-free images. This approach is not limited to label-free imaging such as quantitative phase imaging but extends to virtually any imaging method – whether conventional fluorescence microscopy, label-free techniques, or virtual staining approaches – that can produce intensity maps highlighting cellular structures such as nuclei or membranes. By leveraging these intensity-based representations and employing direct tracking, Ultrack offers an adaptable and efficient solution that bypasses the need for extensive annotation or fine-tuning of segmentation models. Ultrack also excels in processing multichannel data by combining multiple single-channel labels, a feature that, when paired with technologies for engineering multi-colored cells,^50, 51^ promises to enable a new generation of high-accuracy lineage reconstruction. Finally, Ultrack addresses challenges associated with low temporal resolution through GPU-accelerated flow-field registration between frames, broadening its applicability to more experiment designs.

The introduction of a dual-channel sparse labeling approach for validation offers a more challenging and heterogeneous benchmark compared to previous studies focused on specific cell subsets or biological contexts.^15, 16, 31, 64^ This approach paves the way for more robust validation of tracking algorithms over extended periods in large datasets.

Ultrack’s scalability, tested on both high-performance computing clusters and laptops, addresses the growing demand for tools capable of handling increasingly large and complex datasets produced by advanced imaging technologies.^1, 6, 7^ This flexibility in computational requirements democratizes access to high-quality cell tracking across diverse research settings.

Ultrack’s integration with popular platforms like Fiji^35^ and compatibility with established tracking frameworks such as TrackMate^37^ and Mastodon^38^ positions it for widespread adoption. Its multiple interfaces, ranging from graphical user interfaces to command-line tools for high-performance computing environments, align with community efforts to make complex computational tools more accessible to biologists.^19, 20, 33, 34, 36, 79^

Despite all of this, Ultrack has its limitations, as the segmentation hypotheses are only as good as the provided foreground and contour maps. Therefore, despite its robustness to spurious candidate segmentations, when provided with an overwhelming number of incorrect segmentations or systematic errors that are consistent over time, the ILP optimization might favor erroneous segments.

Looking ahead, several promising avenues for future development emerge: (i) Integration of advanced deeplearning techniques, particularly transformer-based models for sequence prediction,^80^ could further improve long-term tracking accuracy. (ii) Expansion of the sparse labeling approach to generate large-scale, high-quality datasets for training and benchmarking future tracking algorithms. (iii) Development of interactive correction tools leveraging Ultrack’s multiple segmentation hypotheses to facilitate efficient manual curation of tracking results. (iv) Exploration of Ultrack’s potential applications beyond developmental biology, including cancer research, immunology, and regenerative medicine.

In conclusion, Ultrack’s innovative approach for joint segmentation and tracking, combined with its scalability and accessibility, has the potential to accelerate research across a wide range of biological disciplines by enabling more accurate, efficient, and accessible cell tracking. Ultrack opens new avenues for investigating complex cellular behaviors, tissue dynamics, and developmental processes at unprecedented scales and resolutions.

## Supplementary Videos

1. Ultrack intuition and overview;
2. Using multiple segmentation hypotheses with temporal consistency helps reduce the challenges of parameter tuning;
3. Enhanced multi-color cell tracking by integrating segmentation results from multiple algorithms;
4. Improved cell tracking in label-free imaging by leveraging virtual staining;
5. Enhancing tracking accuracy through temporal registration;
6. Visualizing Ultrack’s Cell Tracking Challenge benchmark results;
7. Sparse fluorescence labeling enables high-fidelity tracking validation over extended time-lapses;
8. Multi-terabyte cell tracking of zebrafish embryo;
9. Near-perfect nuclearand membrane-based 3D tracking of zebrafish neuromast cells;
10. Ultrack’s guided user interfaces across multiple platforms facilitate cell tracking, catering to diverse user needs and computational environments.

## Code and data availability

The repository for the Ultrack python package and its updated versions are available at https://github.com/royerlab/ultrack. The Fiji plugin can be found at https://github.com/royerlab/ultrack-imagej. The imaging data is accessible at https://public.czbiohub.org/royerlab/ultrack and https://public.czbiohub.org/royerlab/zebrahub/imaging/single-objective

## METHODS

### Animals

All zebrafish care and experimental procedures complied with protocols approved by the Institutional Animal Care and Use Committee at the University of California San Francisco (UCSF). Adult fish were maintained at 28.5 °C and fed twice daily using an automatic feeder.^81^ Embryos were raised at 28.5°C, and staged based on hours post fertilization (hpf). For light-sheet microscopy and injection experiments, we used two transgenic lines: tg(h2afva:h2afva-mCherry) (a gift from J. Huisken, University of Göttingen) and tg(eef1a1/1:H2b-mNeonGreen) (a gift from D. Wagner, UCSF). Zebrafish neuromasts were imaged using the transgenic line tg(cldnb:lyn-mscarlet) x tg(she:h2b-EGFP).^82, 83^

### Multi-colored cell line generation and imaging

MDAMB-231 metastatic breast adenocarcinoma cells (ATCC) were maintained in Dulbecco’s Modified Eagle Medium (DMEM, VWR) with 10% (v/v) fetal bovine serum (FBS, Seradigm VWR), and penicillin and streptomycin (50 U/mL, ThermoFisher Scientific). Cells were stably modified with a variety of RGB-markings with lentiviral gene ontology (LeGO) vector system, as described in Weber *et al*..^52^ Lentivirus was prepared as described in Shah *et al*..^84^ In brief, pseudoviral particles were generated using 293TN cells (System Biosciences, SBI) co-transfected with lentiviral packaging and envelope plasmids (psPAX and pMD2.G, gifts from Didier Trono), and one plasmid of interest containing fluorophore eGFP, mCherry, or EBFP2 (Addgene, #25917, #27339, #85213). Supernatants containing lentivirus were collected, passed through a 0.45 µm filter, concentrated by ultracentrifugation at 200,000 × g for 4 hours at 4°C, resuspended in phosphate-buffered saline (PBS, VWR), and stored at -80 °C. MDA-MB-231s were transduced simultaneously with all three constructs at equal viral titer and a range of multiplicity of infections (MOI). To optimize color diversity, a fraction of cells expressing fluorophore was assessed at five days posttransduction using fluorescent microscopy, and a condition with ≈50% transduction rate of all fluorophores (MOI ≈ 0.7) was selected for subsequent experiments. Cells were seeded 12 h before imaging at a density of ≈ 10,000/well on a glass-bottom plate coated with 50 µg/mL rat tail type I collagen (Corning) in 0.02N glacial acetic acid (Sigma) overnight at 4°C. Imaging was performed on a Keyence BZX800 imaging system equipped with temperature, humidity, and CO2 control, and images were acquired at 10X magnification at ten-minute intervals for 2 days.

### Microinjection for zebrafish sparse labeling

DNA plasmids (pMTB-ef1-H2B-mCherry2 gifted by D. Wagner, UCSF) were extracted and purified with a QIAprep Spin Miniprep Kit (Cat. No. 27104). Purified DNA was diluted to 100 ng/l in water (Invitrogen™ UltraPure™ DNase/RNase-Free Distilled Water, Cat. No. 10977015) and saved in -20. Zebrafish Tol2 mRNA was prepared using pT3TS-Tol2 plasmids shared by Ekker lab.^85^ pT3TS-Tol2 plasmid DNA was linearized by the restriction enzyme BamHI digestion. Linearized plasmids were gel-purified with a QIAquick Gel Extraction Kit (Cat. No. 28704). The 5’-capped mRNAs were synthesized using Invitrogen™ mMESSAGE mMACHINE™ T3 Transcription Kit and purified by lithium included in the kit (ThermoFisher, Cat. No. AM1348). The purified RNA was dried and dissolved with 20 l DNase/RNase-Free water. Dilute it to 200 ng/l and save it as stock in -70 degrees. Before the experiment, the stocked DNA and RNA were thawed, and 1 µL DNA plus 1 µL RNA was mixed with 3 µL of DNase/RNase-Free water containing 0.05% phenyl red. For each microinjection, a 4 nL mixture was injected into a single cell of a 4-8 cell stage embryo from the Tg [ef1-H2B-mNeonGreen] strain. All injections were done with an injector (WPI PV830 Pneumatic Pico Pump), and a micromanipulator (Narishige, Tokyo, Japan). 50 to 100 embryos were injected for each experiment. The injected embryos are ready for mounting and imaging at approximately 4.5 hpf.

### Sample preparation for single objective light-sheet microscopy (DaXi)

Embryos are first dechorionated and then gently embedded into a 1% solution of low-gelling-temperature agarose (Sigma, A0701). The samples are then transferred in a glass-bottom cell culture dish (35mm Cell Culture Dish with Glass Bottom 20mm – Stellar Scientific) and positioned at the correct imaging angle using a custom-made capillary. When solid, the agarose surrounding the tail was cut off and removed using a dissection knife and forceps to permit full development and tail elongation. During the time lapses, the embryos are placed in an embryo-medium solution with 0.016% tricaine to prevent embryo movement.

### Quantitative phase imaging and virtual staining

The A549 cells were cultured at 37°C and 5% CO2 and maintained between 20% and 90% confluency. The cells were seeded on a 12-well glass bottom plate (Cellvis, P12-1.5HN) and imaged using a wide-field microscope (Leica Dmi8). The quantitative phase volumes are virtually stained using the VSCyto2D model to highlight landmark channel organelles such as the nuclei and cell plasma.^57^

### Simultaneous sparse and ubiquitous labels imaging

Four hours post-microinjection, injected embryos (4 hpf) were screened using a Thunder Imager Model Organism (Leica Microsystems). An embryo with 20%-30% H2B-mCherry2 expressing cells (compared to ubiquitous H2B-mNeonGreen cells) was selected and transferred to a glass-bottom petri dish (35mm Cell Culture Dish with Glass Bottom 20mm, Stellar Scientific) containing 0.3x Danieau’s embryo medium. The embryo’s chorion was removed using sharp forceps under the same microscope. The dechorionated embryo was then gently transferred to a petri dish containing 0.1% low-temperature melting agarose (Sigma, A0701) using a glass pipette. Subsequently, the embryo was transferred to FEP tubes (Valco-TFEP130, OD 1/16”, ID 0.03”, Vici Metronics) with agarose via micropipette through a 200 µl tip (Eppendorf). Both ends of the FEP tube were sealed with 2% low-temperature melting agarose to secure the embryo. The tube was placed in the chamber of an adaptive multiview multi-color light-sheet platform^9^ using a customized holder insertion tool filled with low-toxicity silicone (KwikSil™, World Precision Instrument). The mounting system was similar to that previously described in Zebrahub,^9^ but with an upside-down mounting direction. The chamber was filled with embryo water, which was circulated with 0.016% tricaine during imaging. For whole-embryo imaging, the multi-color-interleaved mode was used with an exposure time of 20 ms per channel with the OpenSimView^9^ microscope. The imaging volume comprised 493 z-planes with 1.625 µm z-steps. Images were captured at 90-second intervals, starting at the shield stage (6 hpf), and ending after 24 hpf.

### DaXi microscope parameters

We used our DaXi imaging platform^6^ to perform high-resolution tail development imaging (1.0 NA). We use two orthogonal, oblique light sheets for each time point to improve the image field of view coverage. During imaging, the exposure was set to 20ms, the z-step to 1.24ms, the imaging volume was adapted to each sample, and the time step was around 60 seconds. Time-lapse sessions are about 10 hours long and usually cover a developmental window from 1 or 2 somites to 27 somites.

### Software

The Ultrack package, including its REST API, and additional image analysis routine and plotting were implemented using Blosc, CuPy, FastAPI, imagecodecs, iohub, httpx, ggplot2, napari, networkx, Numba, NumPy, Pandas, PyTorch, scikit-image, SciPy, SLURM, tifffile, traccuracy, uvicorn, websockets, and zarr.^36, 63, 72, 77, 78, 86–92^

### Datasets

Table 2 presents the datasets evaluated in this paper. While not a requirement, some images of the Cell Tracking Challenge were upsampled for improved accuracy.

**Table 2.**
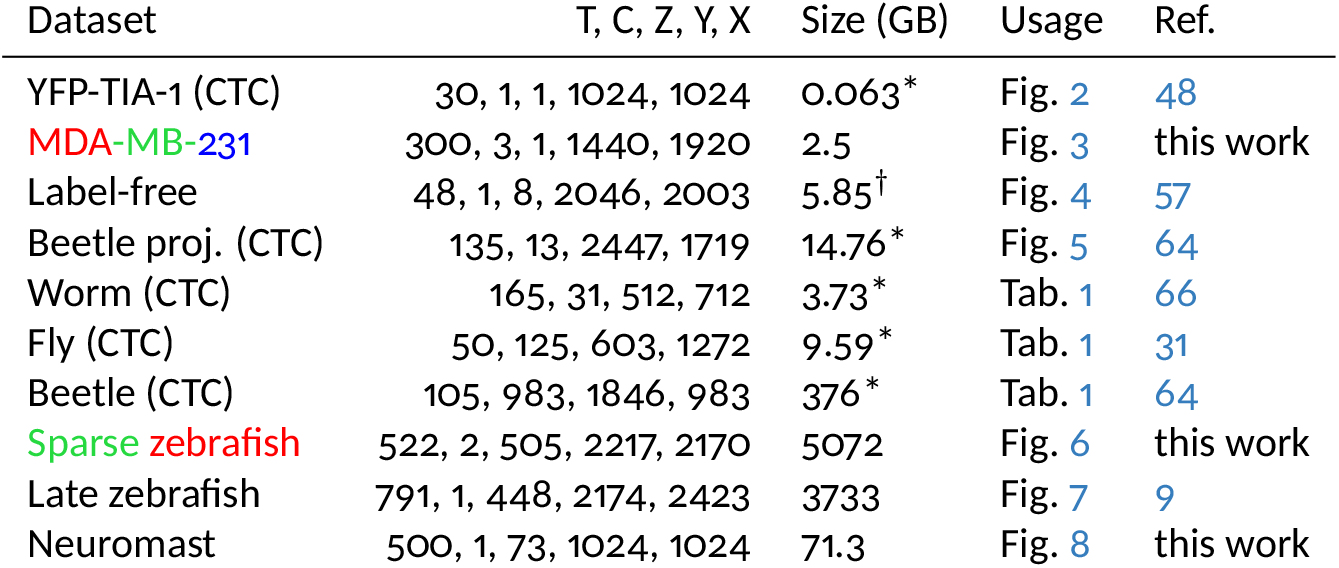
List and references of datasets. †: size before 3D to 2D virtual staining. * indicates averaged size and shape. The size presented in Fig 1f is regarding the input contour map, which can include projections and upsampling.

### Light-sheet imaging data pre-processing

Upon acquisition, the imaging data are converted to the ome-zarr format^93, 94^ using the iohub^87^ Python package. Next, the raw multi-view images are fused. Phase-cross correlation techniques are applied to correct inter-frame movements to ensure stability throughout the time-lapse sequence. In cases where deep-learning methodologies are utilized, image normalization is performed, scaling pixel intensities to the range between 0 and 1 using a user-specified lower and upper quantile, and their background is subtracted using area white top-hat transform. These pre-processing routines and other image processing operations are implemented and accessible via https://github.com/royerlab/dexpv2.

### GPU-accelerated flow field estimation

Flow field estimation of large 3D volumes can be extremely challenging and computationally intensive, often following particle image velocimetry strategies that estimate local translation using phase cross-correlation for small patches of the data.^61^ Instead of maximizing cross-correlation, which minimizes the mean-squared error between frames given a field in the coordinate space, we employ gradient descent to optimize the coordinate space directly. This choice, justified by its enhanced computational efficiency and simplicity, allows for the integration of any differentiable function within the coordinate space. The loss function is defined as:

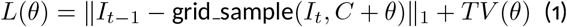

where *I*_*t*_ denotes the frame at time *t, C* represents the identity coordinates (*i.e*. no movement), *θ* indicates the movement flow field we optimize, and *TV* denotes the L1 total variation loss on the flow field. The grid sample function, a differentiable data sampling method from PyTorch,^63^ enables this process. This routine is tailored for specialized GPU hardware, greatly enhancing processing speed. The *θ* optimization is performed for a fixed number of iterations over multiple image resolutions, starting from the lowest resolution, which is then used to initialize the next *θ* optimization stage.

### Label conversion to contour and foreground map

The conversion of multiple segmentation labels into foreground and contour maps involves two main processes. For the foreground map generation: (i) each label is transformed into a binary map, with non-zero pixels assigned a value of one (foreground) and zero pixels remaining as background; (ii) these individual maps are merged into a single foreground map by taking the maximum value across corresponding pixels from all segmentation maps; (iii) this logical OR operation ensures that any pixel belonging to at least one segment is included in the final foreground map. For the contour map generation: (i) to preserve each label and its dual contour, we process each label map to create a binary contour map, where pixels are set to one if their neighboring pixels have differing labels, indicating a boundary; (ii) the final contour map is then created by averaging these individual binary contour maps; (iii) this averaging approach, as opposed to summation, keeps pixel values within the 0 to 1 range.

### Candidate hierarchical (nested) segmentation

The goal of hierarchical segmentation is to compute a set of nested partitions (*i.e*. segmentations) from a multilevel contour map, where each level corresponds to a different segmentation threshold. While a naive approach resorts to binarizing the contour map at each level and computing the connected components of the binary images, Ultrack employs a more sophisticated and computationally efficient method: the hierarchical watershed algorithm.^43, 95^ This algorithm represents the image as a graph *G* = (*V, E*), where *V* are the pixels or voxels, and *E* denotes the edges connecting adjacent voxels. It defines a non-decreasing order of edge weights:

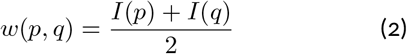

where *p, q* ∈ *E* and *I*(*p*) is the image intensity at a pixel *p* of a contour map. The algorithm then recovers nested segmentations using a region merging approach based on Kruskal’s minimum-spanning tree^96^ and Tarjan’s union-find algorithms.^97^ The process begins with an initial segmentation where each pixel forms its own segment. It then iteratively: (i) selects the minimum unseen edge (*p, q*); (ii) merges the segments of *p* and *q* if they are not already in the same segment; (iii) continues until all edges have been processed. Importantly, this segmentation process operates within each connected component of the binary foreground mask, allowing the algorithm to ignore background regions and thereby greatly speeding up the process. For further details on hierarchical segmentation processing and implementation, see.^45^

### Candidate segmentation association

For each candidate segmentation, we compute the potential temporal associations (*i.e*., tracks). Let ℋ represent the set of all segmentations derived from hierarchical segmentations. We define *E*^*T*^ as the set of pairs representing potential associations between these segmentations, with *w*(·, ·) denoting their respective weights. The association process involves the following steps: (i) For each segmentation *i* ∈ *ℋ*, where *t* denotes the time point of segment *i*, we compute its *k*-nearest neighbors within a predefined radius using their centroid coordinates at time *t* − 1; (ii) These pairs are then included in *E*^*T*^ ; (iii) The association score between any two segmentations *i* and *j* is calculated as *w*(*i, j*) = IoU(*i, j*)^*γ*^. Where *γ* is consistently set to four throughout our study and IoU represents the intersection over union between the segmentation masks of *i* and *j*.

### Segmentation selection and tracking ILP

Given ℋ and *E*^*T*^, as defined previously, we compute the joint segmentation and tracking by solving an integer linear programming (ILP) optimization problem. This ILP framework aims to find the optimal pairing between segments that maximizes their association while adhering to biological constraints. The ILP is formulated as follows:

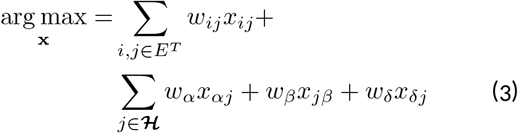

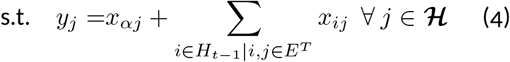

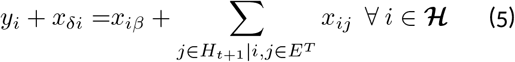

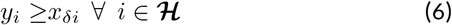

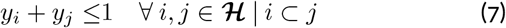

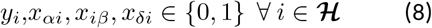

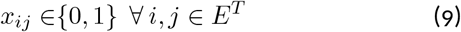

In this formulation, *y*_*i*_ indicates whether a segment is selected, and *x*_*ij*_ indicates the selected associations, *x*_*αi*_, *x*_*iβ*_ and *x*_*δi*_ are slack variables indicating an appearing, disappearing and dividing cell, respectively. Eq. 3 is the objective function to maximize the association between segmentations. The first term in the objective function represents the weight of segmentation associations, whereas the subsequent terms account for appearance, disappearance, and division events, which are user-defined non-positive penalization weights. Eq. 4 and Eq. 5 ensure flow conservation, stipulating that an appearance, division, or incoming association must correspond to at least an outgoing association or a cell disappearance. Eq. 6 restricts cell division to pre-existing cells, and Eq. 7 prevents selecting overlapping segmentations, thereby enforcing that each pixel is uniquely selected.

### Tracking evaluation metrics

The evaluation scores were calculated using official binaries from the Cell Tracking Challenge (CTC) for datasets submitted to the competition. Additional calculations were performed using the traccuracy Python package.^98^ The TRA metric quantifies the cost-effectiveness of editing a predicted lineage into the reference lineage, divided by the cost of constructing the lineage from scratch, *AOGM*_0_. This is mathematically defined as:

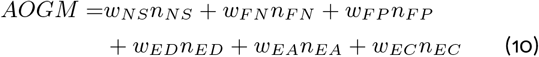

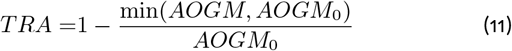

where *n*_*NS*_ represents the number of node splits (*i.e*. undersegmented cells), *n*_*F N*_ and *n*_*F P*_ are false negative and false positive nodes respectively, *n*_*ED*_ and *n*_*EA*_ denote false positive and negative edges, and *n*_*EC*_ represents nodes with incorrect semantics. The weights are set as follows: *w*_*NS*_ = 5, *w*_*F N*_ = 10, *w*_*F P*_ = 1, *w*_*ED*_ = 1, *w*_*EA*_ = 1.5, and *w*_*EC*_ = 1, emphasizing the greater complexity in correcting missing units compared to deleting extra. For further details, refer to Matula et al.^49^ The combined CTC score averages the TRA and SEG scores. The SEG score calculates the average intersection over union (Jaccard index) across all segmentation instances. A ground-truth (*gt*) segmentation instance, *G*_*j*_, is paired to a predicted instance *P*_*i*_ if:

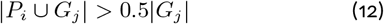

This criterion ensures that the overlap between the predicted and *gt* instances is at least half the size of the *gt* instance, guaranteeing a single match per *gt* instance. The same criterion matches nodes in the *AOGM* metric calculation.

On the multi-color and label-free cell culture experiments where not all lineages are annotated, we applied the recommended CTC protocol, where the first frame of the ground truth is provided to indicate which lineages should be kept, all the remaining lineages are removed, and the CTC metric is computed between this subset of lineages and the annotated data.

### Zebrafish sparse tracking evaluation metrics

The sparse tracking data evaluation uses implementation from the Python package linajea.^24^ The metrics are the proportion of different types of mistakes over the total number of edges (*i.e*. connections between cells of adjacent frames) of the ground-truth data. Because the data is sparse, only a subset of metrics are valid estimates for the remaining unlabeled data, them being:

- FN: Proportion of false negative (*i.e*. missing edges);
- IS: Proportion of identity switches (*i.e*. edges associating incorrect cells);
- FP-D: Proportion of false positive divisions (*i.e*. fake division) within a 1-frame tolerance.
- FN-D: Proportion of false negative divisions (*i.e*. missing division) within a 1-frame tolerance.
- sum: the sum of all the above.

### Color features association

The image intensity information can be used to filter out segments between adjacent frames that are unlikely to represent the same cell. To do that, we compute a normalized distance between the average intensities of segments; if this distance exceeds a predefined threshold *z*, their association is excluded from the tracking Integer Linear Programming (ILP) formulation. Specifically, the relationship must satisfy the following conditions:

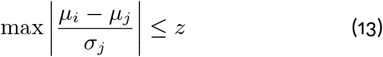

where *µ*_*j*_ is the mean intensity vector within the mask of a segment *S*_*j*_ at time *t* and *σ*_*j*_ the standard deviation, the operations are done element-wise and max results in a scalar from it. Here, compared to a neighboring segment *S*_*i*_ at *t* − 1. The threshold *z* was set to 3.0 throughout the multi-color experiments.

### Fiji plugin and Python interoperability

Fiji^35^ is a Java-based software that can be extended through plugins developed in the same language. To integrate Ultrack with Fiji, we developed a RESTful API^99^ using the FastAPI framework,^88^ facilitating communication between Java and Python. This API employs websockets^100^ for persistent communication, enabling real-time event logging and output streaming from Ultrack to Fiji. We created a graphical user interface (GUI) in Java that serves as a Fiji plugin, leveraging the API provided with the Ultrack Python package. This interface grants access to features and functionalities of both systems. The GUI components and user experience are implemented using the Bootstrap 5^101^ frontend framework. To promote broader interoperability, Ultrack adheres to API specifications detailed in our documentation (https://royerlab.github.io/ultrack/rest_api.html), allowing integration with various programming languages beyond Java and Python.

### Napari plugin

Napari^36^ is a Python library for ndimensional image visualization, annotation, and analysis. It features a GUI built with Qt,^102^ which is extensible with Qt primitives in Python. We developed a custom plugin for Napari integration that still scales with large datasets. In addition to the tracking standard workflow, the plugin offers basic preprocessing operations to fit most users’ needs, which can benefit a significant speedup from GPUs, if any are available. Ultrack’s parameters can be tuned through the GUI. The processing runs in the background in a separate thread, and for that reason, the intermediate results can be visualized using the native Napari viewer. By the tracking execution finish, the results can be exported to several formats, such as Trackmate,^37^ NetworkX graph,^77^ and Zarr array.

## Acknowledgements

We extend our gratitude to Dan Wagner and Jan Huisken for generously sharing their zebrafish lines. We thank Ahmed Abbas and Paul Swoboda for their insightful discussions on Integer Linear Programming and optimization. We are grateful to Bruno Moretti, Veronika Pfannenstill, and Jacky Ko for their valuable feedback and for being early adopters of Ultrack. We appreciate Ivo Sbalzarini and Sandy Schmid for their mentorship and constructive feedback on the manuscript. A.X.F. and I.T. were funded by CNPq 304711/2023-3 and FAPESP 2022/16491-2, 2022/07877-4, and 2023/14427-8. J.L. was funded by NIH Awards R01GM137605 and R35GM153257. This work was funded by the Chan Zuckerberg Biohub San Francisco (CZB SF). We are deeply grateful to CZB SF donors Priscilla Chan and Mark Zuckerberg for their generous support, which made this research possible.

## Author Contributions

J.B., M.L., and L.A.R. conceived the research. J.B. wrote the first version of the Ultrack software, performed the computational experiments, and curated the data except for neuromast and label-free images. I.T. wrote the user interfaces, implemented the REST backend, maintained the software, and trained the model for the sparse/dense labeled embryo. X.Z. prepared and imaged the sparse/dense labeled embryo. M.L. prepared and imaged the zebrahub embryo. T.L. prepared and imaged the neuromast. T.H. and A.B. analyzed the neuromast data. A.J. and A.B. curated the neuromast data. E.H. curated and analyzed the label-free data. S.X. provided support for the microscopy. R.A. prepared and imaged the multicolor data. J.B., I.T., and L.A.R. wrote the paper. J.L., S.M., A.X.F., and A.J. provided guidance. L.A.R supervised the research. All authors contributed to editing the manuscript.

## Notes

https://github.com/royerlab/ultrack

https://public.czbiohub.org/royerlab/ultrack

https://public.czbiohub.org/royerlab/zebrahub/imaging/single-objective

### Competing Interest Statement

The authors have declared no competing interest.

